# Genome-wide co-expression distributions as a metric to prioritize genes of functional importance

**DOI:** 10.1101/2020.01.17.910216

**Authors:** Pâmela A. Alexandre, Nicholas J. Hudson, Sigrid A. Lehnert, Marina R.S. Fortes, Marina Naval-Sánchez, Loan T. Nguyen, Laercio R. Porto-Neto, Antonio Reverter

## Abstract

Genome-wide gene expression is routinely used as a tool to gain a systems-level understanding of complex, biological processes. Numerical approaches that have been used to highlight influential genes include abundance, differential expression, differential variation, network connectivity and differential connectivity. Network connectivity tends to be built on a small subset of extremely high co-expression signals that are deemed significant, but this overlooks the vast majority of pairwise signals. Here, we aimed to assess a complementary strategy, namely whether the entire shape of the distribution of genome-wide co-expression values contains a meaningful biological signal that has hitherto remained hidden from view. We have developed a computational pipeline to assign one of 8 distributions (including normal, skewed, bimodal, kurtotic, inverted) to every gene. We then used a hypergeometric enrichment process to determine if particular genes (regulators versus non-regulators) and properties (differentially expressed or not) tend to be associated with particular distributions greater than would be expected by chance. Examination of several distinct data sets spanning 4 species indicates that there is indeed an additional biological signal present in the genome-wide distribution of co-expression values which would be overlooked by currently adopted approaches.

**Author summary:** High-throughput technologies, such as RNA-Seq, enables access to a vast amount of data. Here, we describe a new approach to interrogate these data and extract further information to help researchers to understand complex phenotypes. Our method is based on gene-level co-expression distributions which were compared to eight possible template shapes to group genes with similar behaviours. The method was tested using five different datasets and the consistency of the results indicate it can be used as a complementary strategy to analyse transcriptomic data.

## Introduction

The uncovering of genetic architecture behind complex phenotypes involves analysing a large variety of genes that interact with each other, and with other molecules, to respond to environmental stimuli [1]. Therefore, gene co-expression studies are becoming increasingly popular in the quest of going beyond differential expression (DE) and recovering more of the functional information from relevant tissues [2]. A gene co-expression study requires the computation of the co-expression correlation coefficient between a given gene and all the other genes under scrutiny, potentially numbered into the thousands for a genome-wide investigation. The result is often a large and tangled network of nodes (genes) connected by edges (significant correlations) from which one expects to extract biological insights and identify regulatory genes. However, defining which gene-level features are relevant to the biological question is not a simple task and main strategies to prioritise genes include focusing on hub (highly connected) genes, using the literature/biological criteria, overlaying other information onto the network such as patterns of differential expression and even exploiting variation in expression around the mean [3]. There is merit in developing additional data driven metrics to dissect co-expression networks and prioritize genes.

One system-wide topological feature of those co-expression networks is a structure comprising many nodes with few connections and few central nodes (hubs) with many connections, in a non-random fashion characterized by a scale-free power-law distribution [4]. The implication is that, within the network, different genes present different “behaviours” (i.e. different abilities to influence other molecules in the network), represented by the strength of their correlation coefficients. We have previously concluded that mRNA present in highly inter-connected modules encode proteins collectively involved in tightly regulated processes such as ribosomal biogenesis (Hudson et al., 2009). However, the typical approach of reverse engineering a co-expression network built from significant pairwise correlations actually reflects only a tiny proportion of the total gene-gene interactions potentially at play across the whole system.

With this in mind, we propose that individual genes may possess diagnostic genome-wide distributions of co-expression values that may be similar or different to what one observes when examining all pairs. Furthermore, these individual distributions may respond to environmental condition and/or physiological state, producing different distributions in different biological circumstances. Hudson et al. (2009) demonstrated that these differences certainly exist at a whole system level by plotting frequency distributions of all pairwise correlation coefficients for six transcriptional landscapes of bovine skeletal muscle, considering different breeds, nutrition and physiological states, each one yielding a distinct distribution. Similarly, Remondini et al. (2005) explored the dynamic properties of all pairwise gene correlation distributions, such as skewness, using a time series expression data of rat fibroblast cell line expressing a conditional Myc-estrogen receptor oncoprotein. Their analysis was able to identify the cascade of c-myc-activated genes within the network. Therefore, current evidence supports the concept that correlation dynamics in co-expression networks have biological meaning.

Given previous studies have explored co-expression distributions and differential co-expression at the level of all gene-pairs, the distribution of correlation coefficients from a single gene across all its competing partners has been neglected. In the approach described here, we aim to determine if those genome-wide patterns are the source of untapped biological information. Based on a template of eight possible distribution shapes, from those likely to arise by chance alone (null distributions) to those apparently subject to some poorly understood external forcing (skewed and bimodal), we evaluated datasets of gene co-expression from human, cattle, duck and fly to determine if correlation distributions can be used as a metric to prioritize genes of functional importance. We anticipate this new metric improving our ability to extract further meaningful information from co-expression networks.

## Methods

The central idea of the methodology developed here was to group individual genes based on them sharing the distribution of genome-wide correlation coefficients. For that purpose, eight shapes were used as templates (Fig 1), varying between unimodal, bimodal, skewed or symmetrical, representing different proportions of positive and negative correlations. Shapes 1 and 2 will be often referred to as null distributions; Shapes 3 and 7 as positively skewed distributions; and Shapes 4 and 8 as negatively skewed distributions.

**Fig 1.**
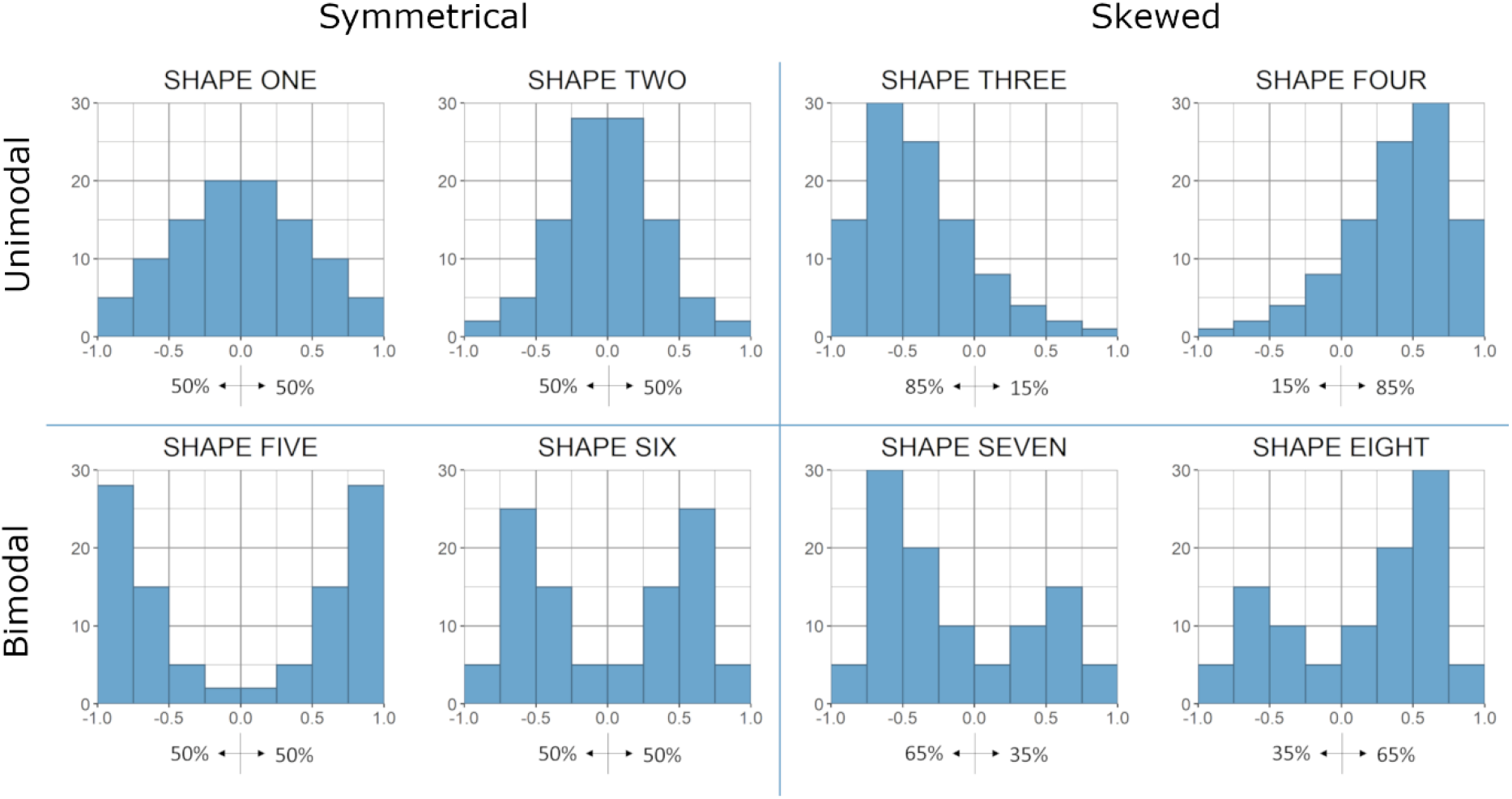
Distribution shapes template. Individual genes were assigned to one of these eight distributions shapes based on their correlation coefficients to all other expressed genes. Distributions were based on the proportion of correlations falling in each of the eight 0.25-bins of the plots.

Starting with a log2 normalized expression matrix, the Pearson correlation coefficient was computed for each gene pair across the samples. For each gene, the number of correlations fell within the eight 0.25-bins of the distributions was recorded (Fig 1) and summary statistics were calculated (i.e.; mean, SD, skewness and kurtosis). Then, based on the proportion of correlations falling in each bin and the summary statistics, and using a naïve chi-square classifier, the Euclidian distance between each gene distribution to each of the eight template distribution shapes was computed. We did this by comparing the observed values for a given gene to the expected values for each template shape listed in Table 1. In algebraical terms, the distance of the *i*-th gene to the *j*-th template shape (*D*_*i,j*_) was computed as follows:

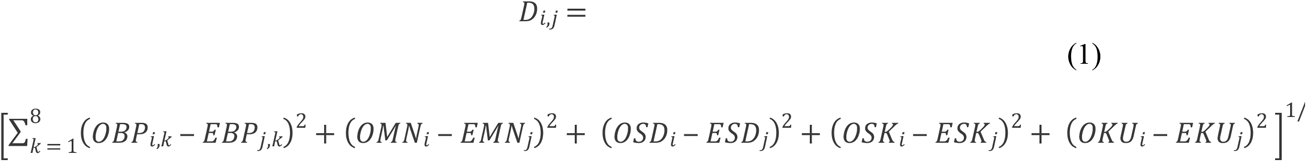

Where subscripts *i*, *j* and *k* indicate gene, template distribution shape and 0.25-bin within shape, respectively; *OBP*_*i,k*_ is the observed bin proportion of the *i*-th gene in the *k*-th bin, that is the proportion of all the co-expression correlation coefficients from the *i*-th gene that fall within the *k*-th 0.25 bin; *EBP*_*j,k*_ is the expected bin proportion of the *k*-th bin in the *j*-th template distribution shape, for instance the expected proportion for the 1^st^ bin in the 1^st^, 2^nd^ and 3^rd^ template shapes are 5%, 2% and 15%, respectively; OMN_*i*_ is the observed mean of all the co-expression correlation coefficients from the *i*-th gene; *EMN*_*j*_ is the expected mean of all the co-expression correlation coefficients in the *j*-th template distribution shape, for instance the expected mean is zero for symmetric distributions such as Shapes 1, 2, 5 and 6; *OSD*_*i*_ is the observed standard deviation of all the co-expression correlation coefficients from the *i*-th gene; *ESD*_*j*_ is the expected standard deviation of all the co-expression correlation coefficients in the *j*-th template distribution shape; *OSK*_*i*_ is the observed skewness of all the co-expression correlation coefficients from the *i*-th gene; *ESK*_*j*_ is the expected skewness of all the co-expression correlation coefficients in *j*-th template distribution shape; *OKU*_*i*_ is the observed kurtosis of all the co-expression correlation coefficients from the *i*-th gene; *EKU*_*j*_ is the expected kurtosis of all the co-expression correlation coefficients in *j*-th template distribution shape.

**Table 1.** Expected bin proportions, mean, standard deviation (SD), skewness and kurtosis for the 8 shapes.

|  | Expected 0.25-Bin Proportions, % |  |  |  |  |  |  |  | Mean | SD | Skewness | Kurtosis |
| --- | --- | --- | --- | --- | --- | --- | --- | --- | --- | --- | --- | --- |
|  | Bin 1 | Bin 2 | Bin 3 | Bin 4 | Bin 5 | Bin 6 | Bin 7 | Bin 8 |  |  |  |  |
| Shape 1 | 5 | 10 | 15 | 20 | 20 | 15 | 10 | 5 | 0 | 0.456 | 0 | -0.660 |
| Shape 2 | 2 | 5 | 15 | 28 | 28 | 15 | 5 | 2 | 0 | 0.354 | 0 | 0.065 |
| Shape 3 | 15 | 30 | 25 | 15 | 9 | 3 | 2 | 1 | -0.387 | 0.382 | 0.857 | 0.640 |
| Shape 4 | 1 | 2 | 3 | 9 | 15 | 25 | 30 | 15 | 0.387 | 0.382 | -0.857 | 0.640 |
| Shape 5 | 28 | 15 | 5 | 2 | 2 | 5 | 15 | 28 | 0 | 0.752 | 0 | -1.771 |
| Shape 6 | 5 | 25 | 15 | 5 | 5 | 15 | 25 | 5 | 0 | 0.566 | 0 | -1.535 |
| Shape 7 | 5 | 30 | 20 | 10 | 5 | 10 | 15 | 5 | -0.137 | 0.532 | 0.5 | -1.111 |
| Shape 8 | 5 | 15 | 10 | 5 | 10 | 20 | 30 | 5 | 0.137 | 0.532 | -0.5 | -1.111 |

Using linear mapping, distances were transformed into similarities (*S*_*i,j*_) of a given gene to belong to each of the template distribution shape as follows:

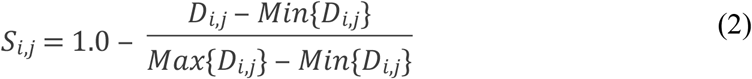

Where *Min*{*D*_*i,j*_} and *Max*{*D*_*i,j*_} are the minimum and maximum *D*_*i,j*_, respectively. Finally, similarities were transformed to probabilities (*P*_*i,j*_) of a given gene to belong to each template distribution shape so that the sum of all probabilities for a given gene added to one:

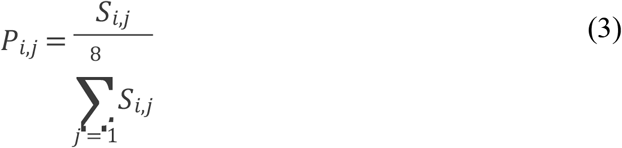

A gene was assigned to the *j*-th distribution template shape if its *P*_*i,j*_ was the largest across all *j*’s. The analyses were implemented using shell scripts to manipulate the data and our own programs in FORTRAN for the fast computation and assignment of within-gene co-expression distributions to template shapes. Scripts and source code are available from the corresponding author upon request.

To gain insight into the biological drivers of gene co-expression distribution, genes were categorized according to their reported biological relevance (e.g.; DE, transcription factors/regulators). A hypergeometric test was applied to identify enriched or depleted categories in each shape, using the function “phyper” in the R environment [7]. Therefore, we compared the within-shape proportion of genes in a given category to the proportion of overall genes in that category. To test the association between the categories and the types of distribution, a chi-square test of independence was applied. Results were considered significant if P-value ≤ 0.05.

To investigate the relationship between number of connections per gene (degree) and distribution shapes, we used the same datasets as input to a PCIT analysis [8]. The PCIT algorithm combines partial correlation coefficients with information theory to determine significant correlations between two genes after accounting for all the other genes under scrutiny. We then computed the number of significant correlations of each gene to evaluate: 1) if there was a relationship between distribution shape and average number of connections per gene using a one-way ANOVA; 2) if the top and the bottom 5% genes based on degree would be enriched in specific shapes.

Finally, although it is not in the scope of this work to discuss specific biological aspects of each dataset, we were interested in evaluating if the different distribution shapes would capture some general biological process. For that purpose, we used the online platform GOrilla (Gene Ontology enRIchment anaLysis and visuaLizAtion tool - http://cbl-gorilla.cs.technion.ac.il/) to tested a list of genes falling in each shape for each dataset against all genes considered for analysis in that dataset. GOrilla uses hypergeometric test and multiple test correction (FDR – false discovery rate) to determine significant enriched gene ontology terms (Padj<0.05). For this analysis, we focused on cell components.

### Datasets used for experimental evaluation

We applied our methodology to publicly available datasets comprised of RNA sequencing (RNA-seq) data from four organisms (cattle, duck, human and fly) in different physiological conditions, from different tissues. These four datasets focussed on different biological questions and were chosen so that we could better explore the utility of new metric. Tables 2 summarizes the characteristics of each dataset.

**Table 2.**
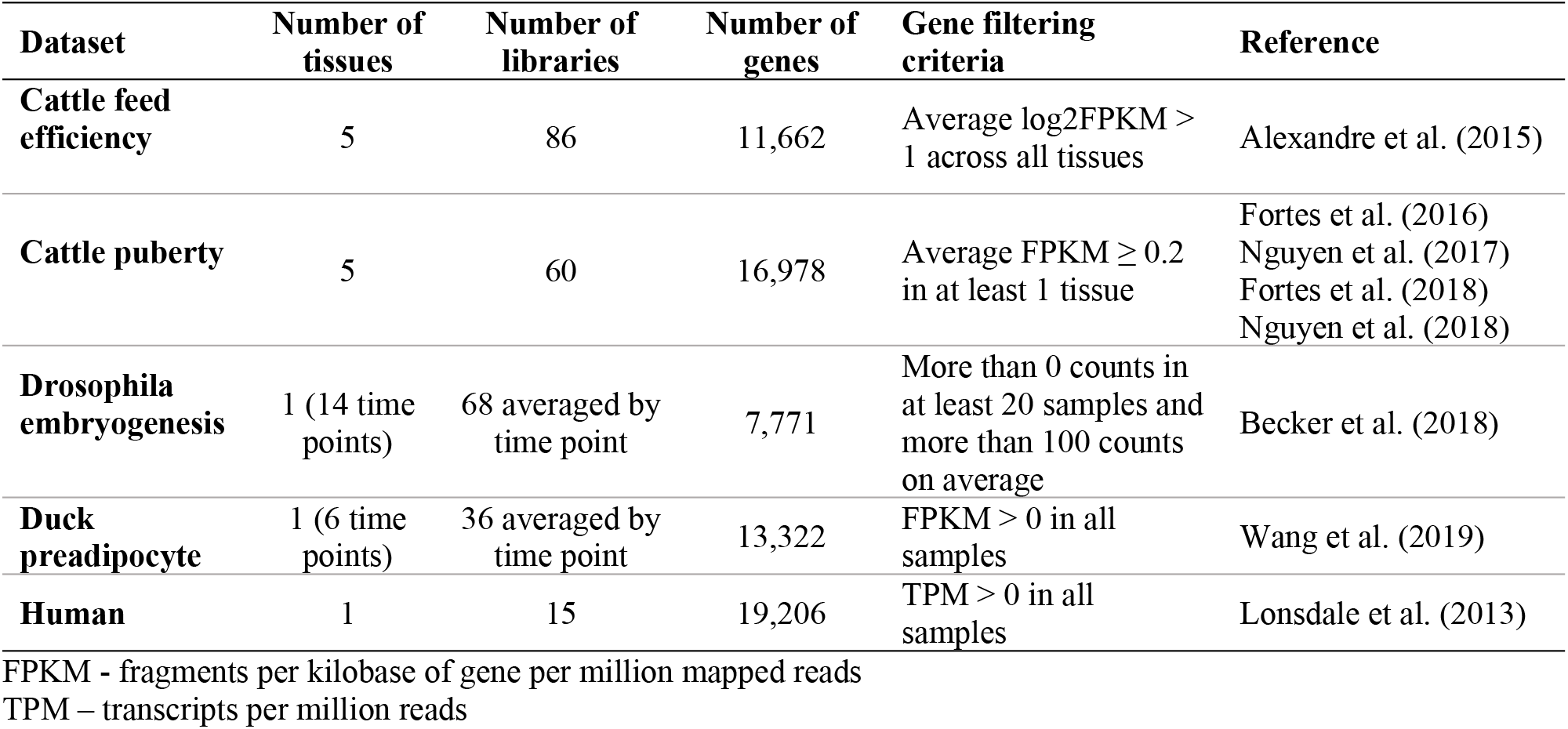
Datasets summary.

| Dataset | Number of tissues | Number of libraries | Number of genes | Gene filtering criteria | Reference |
| --- | --- | --- | --- | --- | --- |
| **Cattle feed efficiency** | 5 | 86 | 11,662 | Average log2FPKM > 1 across all tissues | Alexandre et al. (2015) |
| **Cattle puberty** | 5 | 60 | 16,978 | Average FPKM ≥ 0.2 in at least 1 tissue | Fortes et al. (2016)<br>Nguyen et al. (2017)<br>Fortes et al. (2018)<br>Nguyen et al. (2018) |
| **Drosophila embryogenesis** | 1 (14 time points) | 68 averaged by time point | 7,771 | More than 0 counts in at least 20 samples and more than 100 counts on average | Becker et al. (2018) |
| **Duck preadipocyte** | 1 (6 time points) | 36 averaged by time point | 13,322 | FPKM > 0 in all samples | Wang et al. (2019) |
| **Human** | 1 | 15 | 19,206 | TPM > 0 in all samples | Lonsdale et al. (2013) |

TPM – transcripts per million reads

The cattle feed efficiency dataset is publicly available in the European Nucleotide Archive under the study ID PRJEB27337 (https://www.ebi.ac.uk/ena/data/view/PRJEB27337). Further information about animal management and phenotypic measurements can be found in Alexandre et al. (2015), and information about RNA-Seq libraries and initial processing can be found in Alexandre et al. (2019). Briefly, RNA was extracted from samples of the adrenal gland, hypothalamus, liver, skeletal muscle and pituitary from nine Nellore bulls from each extreme of feed efficiency (evaluated by residual feed intake, Koch et al., 1963). Eighty-six RNA libraries were sequenced on Illumina HiSeq2500 equipment (2×100 pb), reads were aligned to the bovine reference genome (UMD3.1) and expression values were estimated as log2 FPKM (fragments per kilobase of gene per million mapped reads). For this study, we used expression data of 11,662 genes with mean expression > 1 log2(FPKM) across all tissues. Moreover, we classified genes as DE (382 genes) according to Alexandre et al. (2019) and as regulators (REG, 1,072 genes) based on the transcription factors and co-factors indicated in the Animal Transcription Factor Database 2.0 [14].

The cattle puberty dataset is available at EMBL-EBI BioSamples repository (www.ebi.ac.uk/biosamples) under the submission identifiers GSB-113 and GSB-8708. It is comprised of five tissues (hypothalamus, pituitary, ovary, uterus and liver) from 6 pre- and 6 post-pubertal Brahman (*Bos indicus*) heifers, totalizing 60 samples. More details about animal management and data generation can be found in Fortes et al. (2016), Nguyen et al. (2017), Fortes et al. (2018) and Nguyen et al. (2018). In short, libraries were paired-end sequenced using an Illumina HiSeq200 equipment and reads were aligned to the bovine reference genome (UMD3.1). A total of 16,978 genes that presented average FPKM ≥ 0.2 in at least 1 tissue were used for the analysis. Genes were classified as DE (2,335 genes) based on the four aforementioned works and as REG (1,584 genes) also based on the Animal Transcription Factor Database 2.0 [14].

The duck subcutaneous preadipocyte differentiation dataset is available in the Additional File 3 of the source article, where more information about cell culture and RNA-Seq libraries can be found [9]. Briefly, mRNA of six biological replicates of preadipocytes cultured in differentiation medium were collected at −48h, 0h, 12h, 24h, 48h, and 72h and sequenced on an Illumina HiSeq X-10 with PE150. The paired-end reads were aligned to the duck reference genome (Anas_platyrhynchos.BGI_duck_1.0) and gene expression was estimated as FPKM. We kept for analysis only genes presenting counts in all samples (13,322 genes) which were then log2-transformed. Genes were classified as DE (3,321 genes) based on the list provided by the authors in additional file 4 (Wang et al., 2019) and as REG (675 genes) based on the Animal Transcription Factor Database 3.0 [19].

The drosophila embryogenesis dataset is available in NCBI’s Gene Expression Omnibus and is accessible through GEO Series accession numbers GSE121160 and GSE121161. Further information about sample collection and RNA libraries generation can be found in Becker et al. (2018). The mentioned study generated a paired transcriptome/proteome time course dataset with 14 time points during *Drosophila melanogaster* embryogenesis, i.e. 0, 1, 2, 3, 4, 5, 6, 8, 10, 12, 14, 16, 18, 20 hours. The 68 embryo RNA libraries were pooled in equimolar ratio and sequenced on 8 lanes of a HiSeq2500 (1× 51 cycles plus 7 cycles for the index read). Reads were mapped to the BDGP6 fly reference genome and gene expression was estimated as read counts. Data was averaged within each time point and log2 transformed prior to implementation. Genes were classified in the categories defined by Becker et al. (2018), based on pairwise comparison of genes up or down regulated (relative to the first time point - 0h) in mRNA and protein data, resulting in groups named to represent the gene status on mRNA/protein data: up/up (511 genes), down/up (1770 genes), down/down (1048 genes) and up/down (326 genes). Genes were also classified as regulators (791 genes) based on the list provided by Rhee et al. (2014) consisting of essential genes involved in replication and transcription, splicing, DNA repair and cell division.

The human dataset was downloaded from The Genotype-Tissue Expression (GTEx) Project V8 (https://www.gtexportal.org/) which contain data of non-diseased tissue sites across nearly 1000 individuals [10]. We used liver RNA-Seq data from 15 individuals, provided as TPM counts. We kept for analysis only genes presenting non-zero counts in all samples, which were then log2-transformed. Genes were classified as REG (1,153 genes) and tissue enriched (TE, 231 genes) according to information provided by The Human Protein Atlas [22]. Moreover, genes were defined as DE (793 genes) if they were identified by Crow et al. (2019) as having high probability (>0.95) of being DE in any experiment, based on a meta-analysis of over 600 differential expression studies.

## Results and discussion

### Overall co-expression distribution

Although our methodology aims to evaluate co-expression distributions at a gene-level, we did calculate all the correlations across all genes for each of the five RNA-Seq datasets we evaluated: Cattle Feed Efficiency, Cattle Puberty, Drosophila Embryogenesis, Duck Preadipocyte and Human. The overall frequency distributions in each dataset that give us an overview of co-expression patterns can be found in Fig 2. The higher number of positive correlations, even though discrete in some datasets, was already expected and documented on previous research [5,24]. The number of positive correlations is specially elevated in the Cattle Feed Efficiency dataset. This can be due to the high inflammatory response found in the liver of those animals, which remain strong even when analysing all five tissues together and results in a set of highly positively co-expressed genes [12]. On the other side, among the 5 tissues analysed in the Cattle Puberty Dataset, only ovary and uterus showed great differences between pre- and post-puberty and the effect of the coordinated mechanisms regulating those differences are not so strong in the overall frequency distribution. Both Drosophila Embryogenesis and Duck Preadipocyte datasets representing developmental processes through time-series data present similar shapes. They show strong positive and negative correlation as a consequence of the tightly coordinated processes the datasets represent. The Human dataset is the one with the frequency distribution closer to a bell shape, but still, the higher presence of positive correlations can be observed.

**Fig 2.**
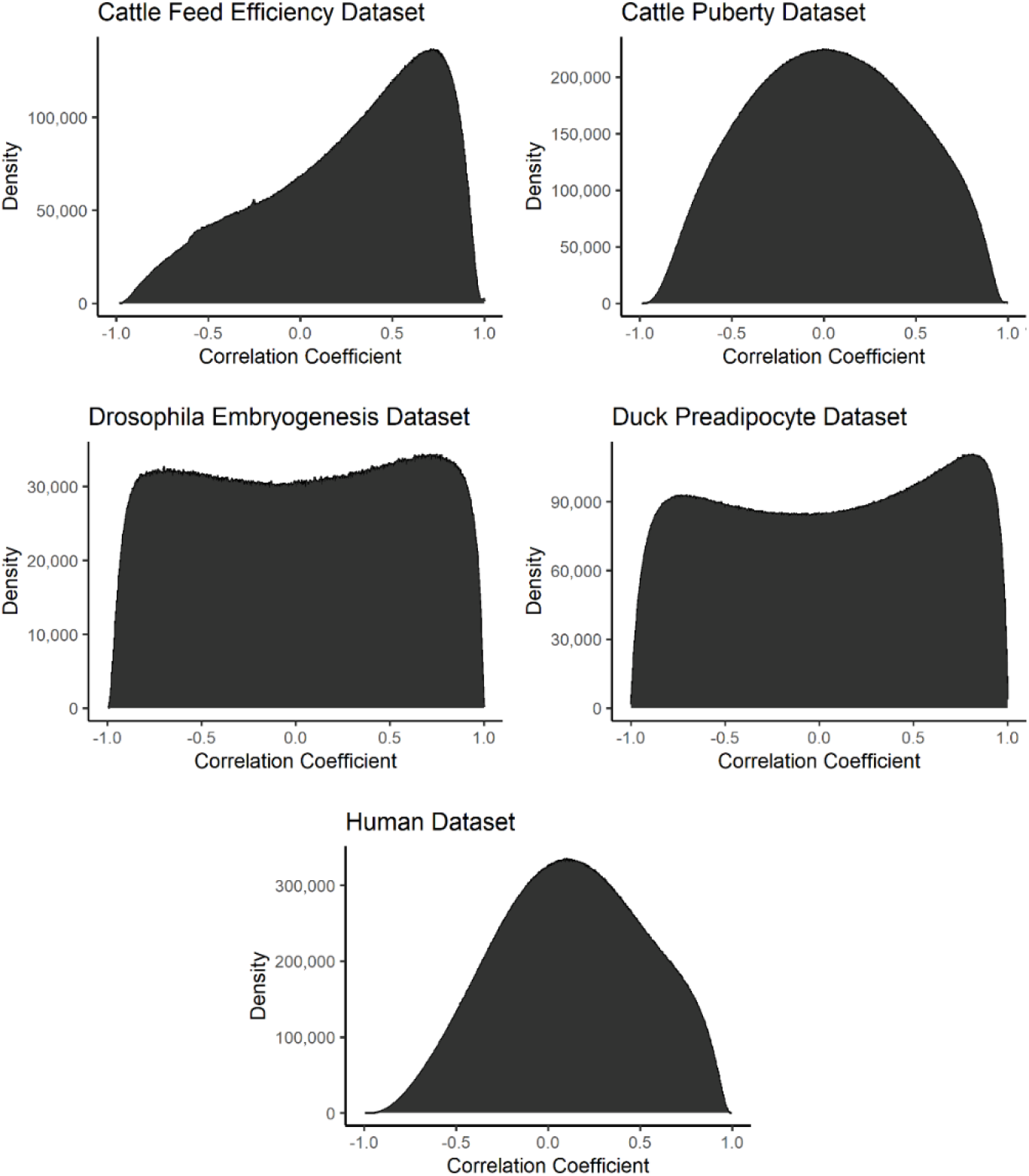
Frequency distributions of all co-expression correlation coefficients in each of the five RNA-Sequence datasets. Datasets include: 1) Cattle Feed Efficiency (11,662 genes and 67,995,291 correlations); 2) Cattle Puberty (16,978 genes and 144,117,753 correlations); 3) Drosophila Embryogenesis (7,771 genes and 30,190,335 correlations); 4) Duck Preadipocyte (13,322 genes and 88,731,181 correlations); and 5) Human (19,311 genes and 186,447,705 correlations).

### Co-expression distribution in datasets with contrasting phenotypes

Considering co-expression distribution at gene-level, the proportion of genes falling in each distribution shape considering the two cattle datasets with contrasting phenotypes can be found in Fig 3. When comparing the proportion of each category of genes with that of all genes within individual distributions, for the cattle feed efficiency dataset we identified an over-representation of REG in negatively skewed distributions (i.e., with an overabundance of positive co-expressions - Shapes 4 and 8) and an under-representation of those genes in null distributions (Shapes 1 and 2) and in a positively skewed distribution (Shape 3). On the other hand, DE genes were under-represented in shape 4 and over-represented in null distributions (Shapes 1 and 2) and in bimodal skewed distributions (Shapes 7 and 8). The exact number of genes falling in each distribution shape and the significance of enrichment analysis can be found in S1 Table.

**Fig 3.**
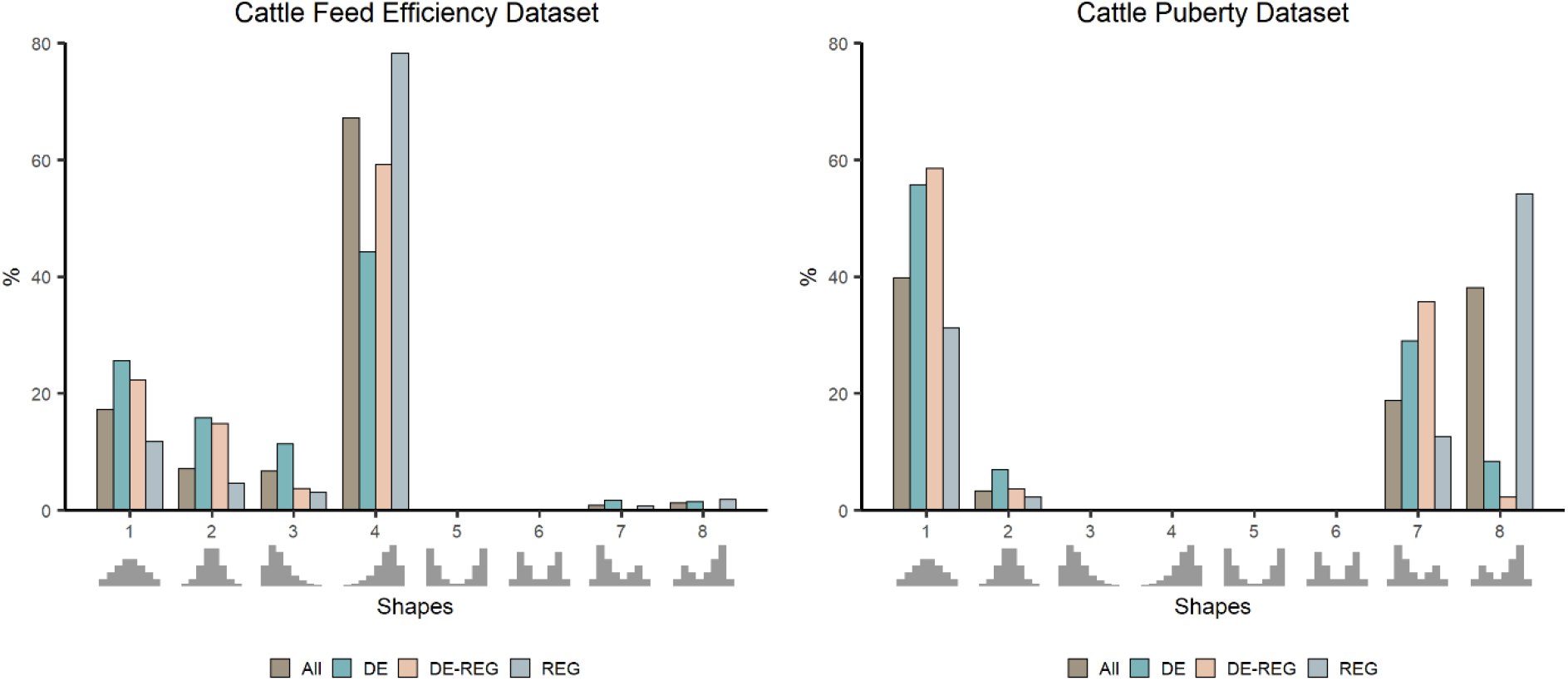
Proportion of genes falling in each template shape in datasets with contrasting phenotypes. The proportion of genes classified as differentially expressed (DE), regulator (REG) or both (DE-REG) are compared to the overall (All) proportion of genes within each shape.

Similarly, the results using the cattle puberty dataset showed an over-representation of REG in a bimodal negative skewed distribution (Shape 8) and an under-representation of those genes in null distributions (Shapes 1 and 2) as well as in a positively skewed distribution (Shape 7). Differentially expressed genes also behave similarly to the previous analysis, being over-represented in null distributions (Shapes 1 and 2) and in a bimodal positively skewed distribution (Shape 7).

Considering that several genes are expected to present different behaviour according to the contrasting condition tested, we applied our pipeline again using the two cattle datasets split by phenotype. We then identified genes that were assigned to different shapes by comparing high to low feed efficiency and pre to post puberty. From the 11,662 genes tested using the cattle feed efficiency dataset, 2,032 genes were assigned to different shapes depending on the condition, being 133 DE and 158 REG. Likewise, 2,740 out of 16,978 genes from the cattle puberty dataset were assigned to different shapes depending on the phenotype, being 620 DE and 216 REG. The shift in the proportion of each class of gene depending on the phenotypic condition can be seen in Fig 4.

**Fig 4.**
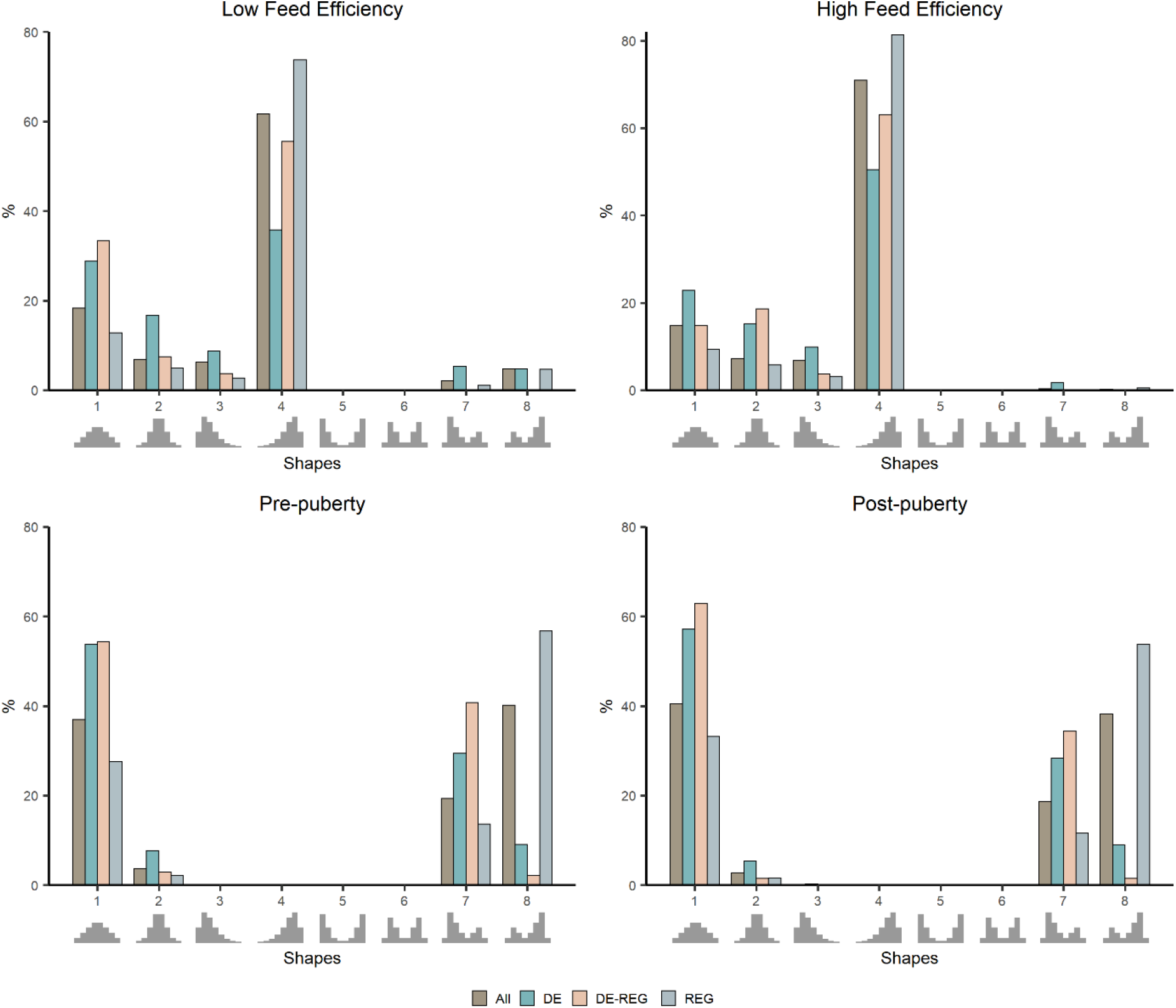
Proportion of genes falling in each template shape in contrasting phenotypic conditions within cattle datasets. The proportion of genes classified as differentially expressed (DE), regulator (REG) or both (DE-REG) are compared to the overall (All) proportion of genes within each shape.

The enrichment analysis for DE and REG among the genes changing behaviour according to the phenotype were again concordant, with REG being under-represented and DE being over-represented (P<0.01) in both datasets. Most REG genes, such as transcription factors and co-factors, are expected to be central genes in the network, presenting co-expression with many genes as a consequence of their regulatory role in highly coordinated biological processes. This can be one reason why they are not particularly enriched among the genes changing co-expression distribution between conditions. Nevertheless, the few REG that do change co-expression distribution between conditions are defiantly worth to be further explored as potential key regulators. On the other hand, a gene identified as DE, being either central to a specific function or the final product of an altered pathway, are more prone to be condition-specific/enriched, not only regarding their expression level but also regarding their relationship with other genes. By identifying DE genes of central or peripheral function in the network based on their enrichment in null or non-null distributions, and exploring changes in their behaviour, one can gain insight into the molecular dynamics behind the phenotype regulation.

In general, genes changing behaviour between conditions might be DE genes that fail the significance threshold in the differential expression analysis. They can also be genes that, although not differentially expressed between conditions, play different roles depending on the overall gene expression pattern, a feature already widely explored using differential connectivity measures [25]. The advantage of comparing distribution shapes is that it considers the direction (positive or negative) of the correlations and the proportion of the correlation falling in each bin of the distribution, corresponding to the correlation’s strength. One can explore all genes changing behaviour or focus on specific distribution changes, for instance, from unimodal to bimodal or from symmetric to skewed. In our datasets, 38% of genes changing shapes in Feed Efficiency and 77% in Puberty represents a change from null to non-null distributions or vice-versa (which, in this case, also represents changes from symmetrical to skewed). Respectively, 37% and 76% genes changing shapes are moving from unimodal to bimodal distributions or vice-versa. Although percentages are similar, different sets of genes are selected depending on the criteria.

The enrichment analysis for DE and REG in each shape show similar results between phenotypic conditions (S2 and S3 Tables). Nevertheless, it is possible to observe a shift in the proportion of genes falling in each shape, particularly from bimodal to unimodal shapes, implicating in a loss/gain of genes presenting both positive and negative correlations at the same time.

### Co-expression distribution in time series datasets

Both time series datasets, duck fat differentiation and drosophila embryogenesis, follow the same pattern identified in the cattle data sets (Fig 5), with REG being under-represented in a null distribution (Shape 1) and over-represented in a bimodal skewed distribution (Shape 7 or 8, S4 Table). Although in the Drosophila dataset the DE genes were subdivided into different classes according to their concordance between mRNA and protein expression data, it is possible to observe in both datasets the enrichment of DE genes being split between null and non-null distributions. Quite remarkably, considering non-null distribution in the Drosophila dataset, genes consistently down-regulated (down-down) are enriched in the negatively-skewed bimodal distribution (Shape 8) while genes consistently up-regulated (up/up) are enriched in the opposite shape - the positively-skewed bimodal distribution (Shape 7). Moreover, those two classes are under-represented in the symmetrical bimodal distribution (Shape 6) where up/down and down/up are both enriched.

**Fig 5.**
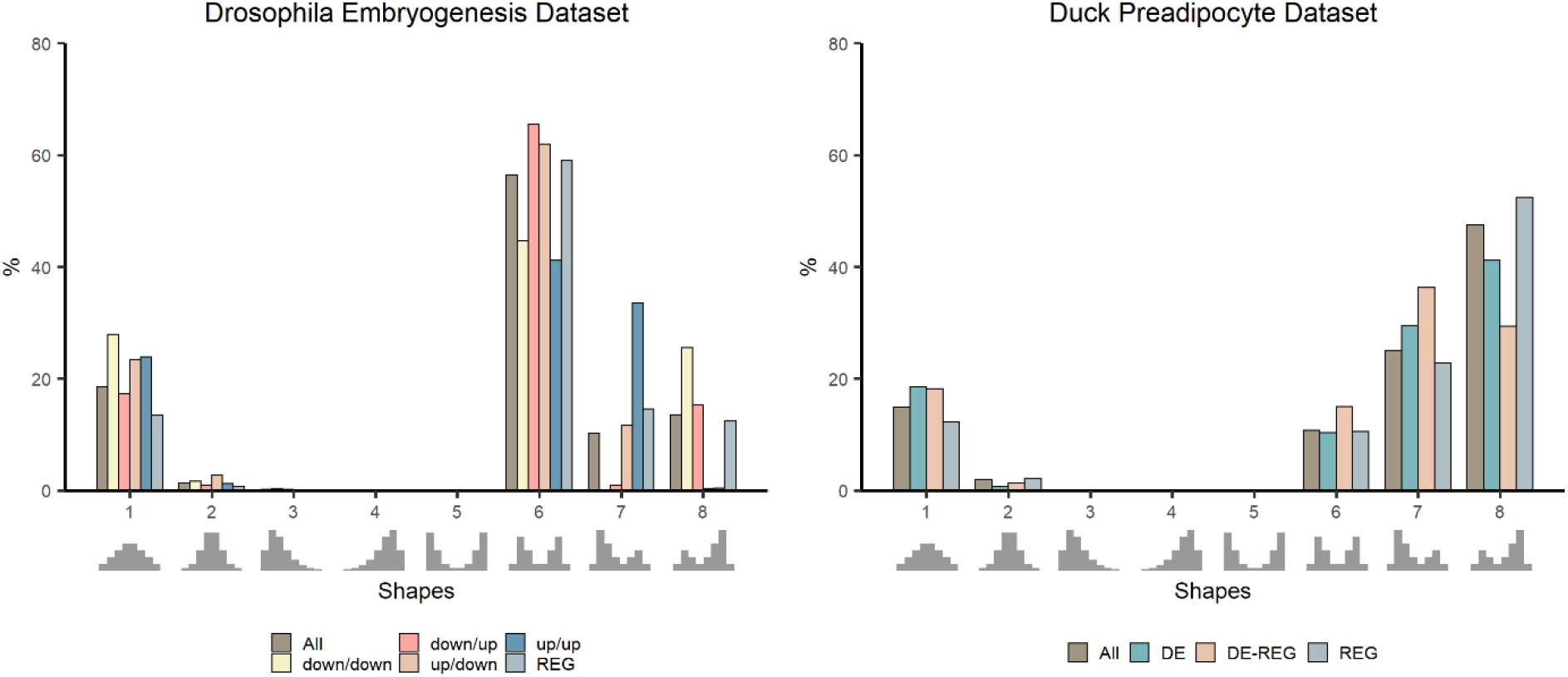
Proportion of genes falling in each template shape in time series datasets. The proportion of genes classified as differentially expressed (DE), regulator (REG) or both (DE-REG) are compared to the overall (All) proportion of genes within each shape. In Drosophila dataset, DE genes were clustered into four groups (down/down, down/up, up/down and up/up) – refer to methods for more information.

Another curious observation is that, while both bovine datasets (representing contrasting phenotypes) present no genes in Shape 6, both time-series datasets not only present gene falling in this shape but also there is an enrichment of DE genes. It is important to notice at this point the impact of species, tissues and even filtering criteria in the proportion of genes assigned to each shape in each dataset. Although some patterns can be identified, each dataset has its idiosyncrasies and different aspects may be worth investigating.

### Co-expression distribution in a physiological baseline dataset

In contrast to the other datasets, the Human dataset consists of several datapoints all representing a single “non-disease” baseline state and this fact is reflected in the results (Fig 6). The REG, found in other datasets to be over-represented in skewed distributions, are over-represented in a null distribution (Shape 2) and under-represented in skewed bimodal distributions (Shapes 7 and 8). In the previous datasets, DE genes already showed enrichment in null distributions, but they also showed enrichment in other non-null shapes. Without the effect of contrasting conditions, genes with high probability of being differentially expressed were enriched in null distributions (Shapes 1 and 2) and in Shape 3 but with very few genes falling in this last shape (43 genes – 0.2% of the total). Tissue enriched genes was the only category enriched in non-null distribution (Shape 7). That is probably because genes particular to tissue’s specific activities tend to be tightly correlated, as they need to be expressed in a coordinated pattern to keep tissues’ functions despite physiological conditions. That behaviour can be clearly observed in studies that constructed co-expression networks using multiple tissue transcriptomic data, where tissue-specific genes push genes to cluster by tissue [12,26].

**Fig 6.**
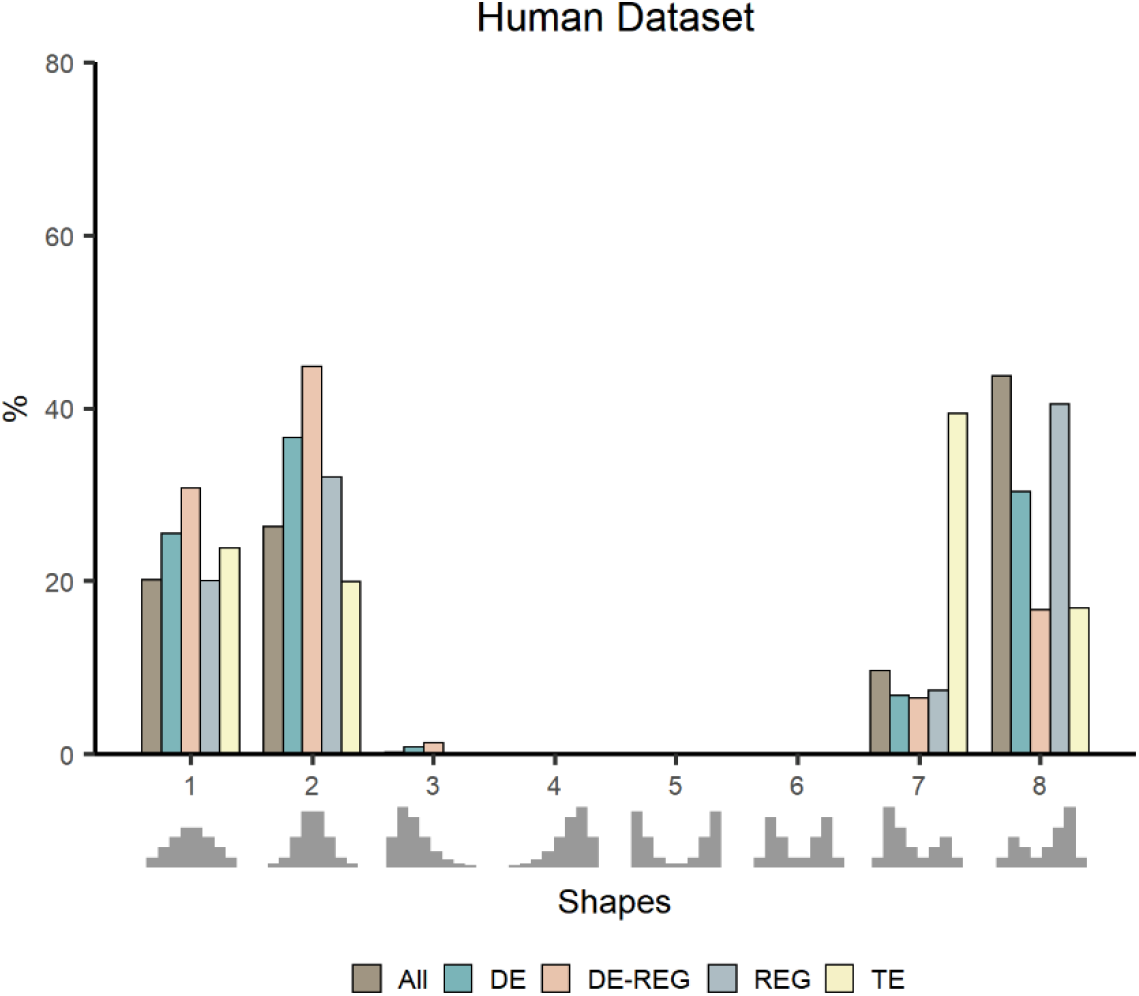
Proportion of genes falling in each template shape in human dataset. The proportion of genes classified as differentially expressed (DE), regulator (REG) or both (DE-REG) are compared to the overall (All) proportion of genes within each shape.

### Relationship between gene categories and distribution shapes

Considering the similarities observed between the first three datasets and the fact that without contrasting phenotypes genes with regulatory potential appear enriched in null distributions (i.e., Shapes 1 and 2), we evaluated the dependence between DE or REG genes and null vs non-null (Shapes 3 to 8), unimodal (Shapes 1 to 4) vs bimodal (Shapes 5 to 8) and symmetric (Shapes 1, 2, 5, and 6) vs skewed (Shapes 3, 4, 7 and 8) distributions. Again, we found a consistent pattern among most of the datasets, with DE genes being found more frequently than expected (P<0.05) in null, unimodal and symmetrical distributions and REG being found more frequently than expected (P<0.05) in non-null, bimodal and skewed distributions (Fig 7). On the other hand, for the human dataset, both DE and REG were found more frequently than expected (P<0.05) in null, unimodal and symmetrical distributions.

**Fig 7.**
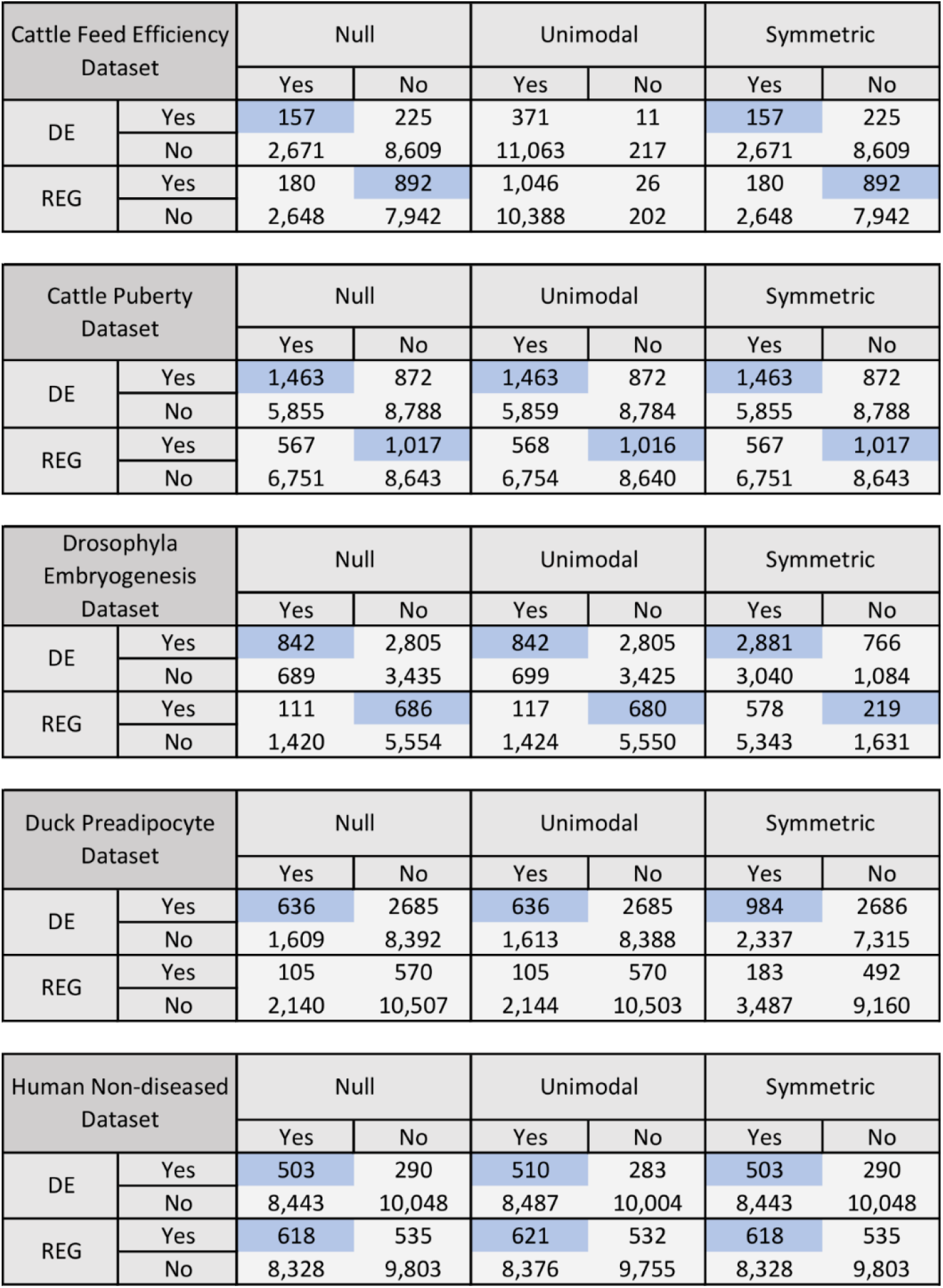
Number of genes assigned to different distribution shapes. More especially, the tables show the number of differentially expressed genes (DE) and regulator (REG) genes assigned to null (Shapes 1 and 2), unimodal (Shapes 1 to 4) and symmetric (Shapes 1, 2, 5 and 6) distributions. Blue cells represent the type of distribution in which DE or REG were found in higher number than expected by chance where variables were found to be dependent (P<0.05).

### Relationship between gene degree and distribution shapes

Because genes that are expected to be highly connected to others (such and REG and TE) appear enriched in non-null distributions, we explored the relationship between number of significant correlations and distributions shapes. Table 3 shows that, for all datasets, there is a significant difference in the average number of connections per gene in each distribution shape.

**Table 3.** Average number of connections per gene (Log10) falling in each shape according to the dataset and P-value for the one-way ANOVA.

| Dataset | Average number of connections per gene (Log10) |  |  |  |  |  |  |  | P-value |
| --- | --- | --- | --- | --- | --- | --- | --- | --- | --- |
|  | Shape 1 | Shape 2 | Shape 3 | Shape 4 | Shape 5 | Shape 6 | Shape 7 | Shape 8 |  |
| **Cattle feed efficiency** | 3.04 | 2.81 | 3.30 | 3.49 | - | - | 3.15 | 3.20 | <2E-16 |
| **Cattle puberty** | 3.14 | 2.52 | 1.40 | 1.87 | - | - | 3.57 | 3.58 | <2E-16 |
| **Drosophila embryogenesis** | 2.58 | 2.26 | 2.39 | - | - | 3.03 | 2.92 | 2.87 | <2E-16 |
| **Duck preadipocyte** | 2.33 | 2.06 | 1.82 | - | - | 2.60 | 2.60 | 2.66 | <2E-16 |
| **Human** | 2.60 | 2.34 | 2.46 | 2.40 | - | - | 3.17 | 3.44 | <2E-16 |

As anticipated, the bottom 5% genes regarding degree (less connected genes) were enriched in both null distributions (Shapes 1 and 2) for all datasets (S6 Table). The few enrichments out of those 2 shapes were due to the low number of genes assigned to that shape. On the other hand, bottom genes were depleted in non-null distributions, particularly in bimodal shapes. The top 5% genes (corresponding to genes with the higher number of significant correlations) were enriched in different shapes depending of the dataset, but exclusively in non-null distribution. Except for the Human dataset, those enrichments were found in shapes that were also enriched for functionally important categories of genes (DE or REG). This overlap, particularly between transcription factors and highly connected genes have been reported before [5]. For all datasets, top genes were depleted in null distributions. This results reinforce that genes related to tightly coordinated processes are more often found in non-null distribution.

### Functional enrichment within distribution shape

The functional enrichment of genes falling in each shape, although not consistently among the different datasets, demonstrate strong biological signals resulting from our methodology (Table 4), with some of the enrichments as significant as q-value of 1.51E-153 (nucleoplasm). To the best of our knowledge, this is the first time it is demonstrated genes grouped according to the distribution of their co-expression correlations can capture specific gene functions. Here, we reported enrichment results at Cellular Component level, as it is not the focus of this work to discuss particular biological mechanisms behind each dataset. Nevertheless, we also have found shape-specific enrichment for Biological Process and Molecular Function in our test datasets (data not shown).

**Table 4.** Hypergeometric gene set enrichments for each distribution within each data set (FDR q-values) based on Cellular Component.

|  | Feed efficiency | Puberty | Drosophila | Duck | Human |
| --- | --- | --- | --- | --- | --- |
| **Shape 1** | Ribosomal unit ($7.84E-15$)<br>Mitochondrial part ($1.61E-14$) | Membrane part ($9.17E-32$) | Cytoplasmic part ($1.67E-8$)<br>Mitochondrion ($1.89E-4$) | Condensed chromatin outer kinetochore ($3.42E-3$) | Plasma membrane part ($1.95E-5$) |
| **Shape 2** | Mitochondrial part ($3.82E-13$)<br>ribosomal unit ($1.09E-3$) | Condensed chromatin outer kinetochore ($3.91E-2$) | No functional enrichment | No functional enrichment | Immunoglobulin complex ($2.98E-21$) |
| **Shape 3** | Contractile fibre ($7.55E-24$) | No functional enrichment | No functional enrichment | No functional enrichment | No functional enrichment |
| **Shape 4** | Nucleoplasm ($2.31E-27$) | No functional enrichment | | | No functional enrichment |
| **Shape 5** | - | - | - | - | - |
| **Shape 6** | - | - | Nuclear part ($7.64E-26$) | Proteasome complex ($6.71E-3$) | - |
| **Shape 7** | No functional enrichment | - | Cytosolic part ($9.75E-27$) | Ribosomal subunit ($5.04E-16$) | Peroxisomal matrix ($3.83E-8$) |
| **Shape 8** | Neuron projection ($2.51E-2$) | Nucleoplasm ($1.51E-153$) | Cytoplasm ($1.45E-6$) | Golgi apparatus ($2.4E-2$) | Intracellular organelle part ($1.29E-108$) |

## Conclusion

We started with the premise that in a co-expression network, different genes present different co-expression distributions and can be grouped according to those. Indeed, considering the five vastly different datasets we analyzed, genes were assigned consistently to pre-defined distribution shapes, regarding the enrichment of differentially expressed and regulatory genes, in situations involving contrasting phenotypes, time series or physiological baseline data.

Admittedly, there is some arbitrary subjectivity in the creation of the proposed 8 template shapes. However, we believe 8 is the minimum number of shapes required to capture in a balanced way the symmetry vs. skewness contrast in one dimension, and the uni- vs. multi-modality contrast in another dimension. Similarly, the use of more or less bins within distributions (eg., having 10 0.10-bins instead of 8 0.25-bins) is worthy of further research. Indeed, across the five datasets analyzed, no gene was allocated to Shape 5.

In conclusion, the results highlight that the distribution shape of correlation coefficients can be used as a novel metric to prioritise genes of functional importance and to further explore topological characteristics of gene networks. By considering that highly connected genes will be assigned to particular distribution shapes according to the experimental design underlying the gene co-expression networks, regulatory genes can even be identified in datasets that do not represent physiological contrasts or time series.

## Acknowledgments

The authors thank Dr. Yutao Li and Dr. Sonja Dominik for reviewing the manuscript.

---

**S1 Table.** Number of genes assigned to each distribution shape in datasets with contrasting phenotypes. Numbers in parentheses represent the P-values for enrichment (above) and depletion (below). The categories include differentially expressed (DE) and regulator (REG) genes.

|  | Cattle Feed Efficiency Dataset |  |  |  |  | Cattle Puberty Dataset |  |  |  |  |
| --- | --- | --- | --- | --- | --- | --- | --- | --- | --- | --- |
|  | REG | DE | DE-REG | Others | Total | REG | DE | DE-REG | Others | Total |
| **Shape 1** | (1)<br>122<br>(1.62E-07) | (2.93E-05)<br>91<br>(1) | (0.31)<br>6<br>(0.83) | (0.03)<br>1,784<br>(0.98) | 2,003 | (1)<br>449<br>(4.87E-13) | (8.70E-59)<br>1,223<br>(1) | (5.14E-06)<br>82<br>(1) | (1)<br>5,003<br>(4.19E-21) | 6,757 |
| **Shape 2** | (1)<br>48<br>(3.61E-04) | (9.20E-09)<br>56<br>(1) | (0.12)<br>4<br>(0.96) | (0.80)<br>717<br>(0.23) | 825 | (1)<br>31<br>(4.39E-03) | (2.11E-20)<br>153<br>(1) | (0.49)<br>5<br>(0.68) | (1)<br>372<br>(1.82E-10) | 561 |
| **Shape 3** | (1)<br>31<br>(2.10E-08) | (0.08)<br>40<br>(0.95) | (0.85)<br>1<br>(0.45) | (3.73E-03)<br>708<br>(1) | 780 | (1)<br>0<br>(0.84) | (1)<br>0<br>(0.76) | (1)<br>0<br>(0.98) | (0.60)<br>2<br>(1) | 2 |
| **Shape 4** | (5.94E-17)<br>818<br>(1) | (1)<br>157<br>(1.94E-19) | (0.86)<br>16<br>(0.25) | (0.98)<br>6,835<br>(0.02) | 7,826 | (0.16)<br>1<br>(0.99) | (1)<br>0<br>(0.76) | (1)<br>0<br>(0.98) | (0.95)<br>1<br>(0.40) | 2 |
| **Shape 5** | (1)<br>0<br>(1) | (1)<br>0<br>(1) | (1)<br>0<br>(1) | (1)<br>0<br>(1) | 0 | (1)<br>0<br>(1) | (1)<br>0<br>(1) | (1)<br>0<br>(1) | (1)<br>0<br>(1) | 0 |
| **Shape 6** | (1)<br>0<br>(1) | (1)<br>0<br>(1) | (1)<br>0<br>(1) | (1)<br>0<br>(1) | 0 | (1)<br>0<br>(1) | (1)<br>0<br>(1) | (1)<br>0<br>(1) | (1)<br>0<br>(1) | 0 |
| **Shape 7** | (0.74)<br>7<br>(0.40) | (7.63E-111)<br>6<br>(1) | (1)<br>0<br>(1) | (0.76)<br>80<br>(0.35) | 93 | (1)<br>182<br>(4.22E-11) | (3.11E-36)<br>636<br>(1) | (1.52E-06)<br>50<br>(1) | (1)<br>2,315<br>(7.78E-14) | 3,183 |
| **Shape 8** | (0.03)<br>19<br>(0.98) | (2.47E-08)<br>5<br>(1) | (1)<br>0<br>(1) | (0.98)<br>111<br>(0.04) | 135 | (5.86E-38)<br>781<br>(1) | (1)<br>183<br>(4.99E-251) | (1)<br>3<br>(5.12E-25) | (1.43E-75)<br>5,506<br>(1) | 6,473 |
| **Total** | 1,045 | 355 | 27 | 10,235 | 11,662 | 346 | 236 | 9 | 1,952 | 16,978 |

**S2 Table.** Number of genes assigned to each distribution shape in the cattle feed efficiency dataset split by phenotype. Numbers in parentheses represent the P-values for enrichment (above) and depletion (below). The categories include differentially expressed (DE) and regulator (REG) genes.

|  | Low Feed Efficiency |  |  |  |  | High Feed Efficiency |  |  |  |  |
| --- | --- | --- | --- | --- | --- | --- | --- | --- | --- | --- |
|  | REG | DE | DE-REG | Others | Total | REG | DE | DE-REG | Others | Total |
| Shape 1 | (1) | (7.67E-07) | (0.05) | (0.12) | 2,137 | (1) | (3.18E-05) | (0.58) | (0.01) | 1,729 |
|  | 134 | 102 | 9 | 1,892 |  | 98 | 81 | 4 | 1,546 |  |
|  | (3.49E-07) | (1) | (0.98) | (0.89) |  | (2.97E-08) | (1) | (0.63) | (0.99) |  |
| Shape 2 | (1) | (1.39E-10) | (0.56) | (0.96) | 800 | (0.97) | (7.37E-08) | (0.04) | (0.97) | 830 |
|  | 52 | 59 | 2 | 687 |  | 60 | 54 | 5 | 711 |  |
|  | (0.01) | (1) | (0.72) | (0.05) |  | (0.04) | (1) | (0.99) | (0.03) |  |
| Shape 3 | (1) | (0.04) | (0.83) | (1.96E-04) | 732 | 1 | 0.01 | 0.85 | 5.89E-04 | 786 |
|  | 28 | 31 | 1 | 672 |  | 32 | 35 | 1 | 718 |  |
|  | (2.14E-08) | (0.98) | (0.49) | (1) |  | (3.66E-08) | (0.99) | (0.45) | (1) |  |
| Shape 4 | (3.75E-18) | (1) | (0.80) | (0.97) | 7,192 | (3.66E-16) | (1) | (0.87) | (0.99) | 8,267 |
|  | 771 | 127 | 15 | 6,279 |  | 850 | 179 | 17 | 7,221 |  |
|  | (1) | (1.16E-23) | (0.32) | (0.03) |  | (1) | (1.42E-16) | (0.24) | (0.02) |  |
| Shape 5 | (1) | (1) | (1) | (1) | 0 | (1) | (1) | (1) | (1) | 0 |
|  | 0 | 0 | 0 | 0 |  | 0 | 0 | 0 | 0 |  |
|  | (1) | (1) | (1) | (1) |  | (1) | (1) | (1) | (1) |  |
| Shape 6 | (1) | (1) | (1) | (1) | 0 | (1) | (1) | (1) | (1) | 0 |
|  | 0 | 0 | 0 | 0 |  | 0 | 0 | 0 | 0 |  |
|  | (1) | (1) | (1) | (1) |  | (1) | (1) | (1) | (1) |  |
| Shape 7 | (1) | (2.08E-04) | (1) | (0.52) | 248 | (1) | (4.21E-04) | (1) | (0.90) | 33 |
|  | 11 | 19 | 0 | 218 |  | 0 | 6 | 0 | 27 |  |
|  | (4.71E-03) | (1) | (0.56) | (0.56) |  | (0.04) | (1) | (0.93) | (0.21) |  |
| Shape 8 | (0.56) | (0.52) | (1) | (0.44) | 553 | (0.01) | (1) | (1) | (0.99) | 17 |
|  | 49 | 17 | 0 | 487 |  | 5 | 0 | 0 | 12 |  |
|  | (0.50) | (0.58) | (0.27) | (0.61) |  | (1) | (0.59) | (0.96) | (0.05) |  |
| Total | 1,045 | 355 | 27 | 10,235 | 11,662 | 1,045 | 355 | 27 | 10,235 | 11,662 |

**S3 Table.** Number of genes assigned to each distribution shape in the cattle puberty dataset split by phenotype. Numbers in parentheses represent the P-values for enrichment (above) and depletion (below). The categories include differentially expressed (DE) and regulator (REG) genes.

|  | Pre-puberty |  |  |  |  | Post-puberty |  |  |  |  |
| --- | --- | --- | --- | --- | --- | --- | --- | --- | --- | --- |
|  | REG | DE | DE-REG | Others | Total | REG | DE | DE-REG | Others | Total |
| **Shape 1** | (1)<br>397<br>(1.61E-15) | (6.66E-66)<br>1,179<br>(1) | (2.05E-05)<br>76<br>(1) | (1)<br>4,620<br>(1.61E-22) | 6,272 | (1)<br>478<br>(1.32E-09) | (4.66E-64)<br>1,254<br>(1) | (6.50E-08)<br>88<br>(1) | (1)<br>5,046<br>(6.50E-28) | 6,866 |
| **Shape 2** | (1)<br>31<br>(5.45E-04) | (9.98E-23)<br>169<br>(1) | (0.75)<br>4<br>(0.42) | (1)<br>412<br>(1.86E-10) | 616 | (1)<br>23<br>(2.81E-03) | (4.12E-14)<br>118<br>(1) | (0.89)<br>2<br>(0.27) | (1)<br>314<br>(3.77E-06) | 457 |
| **Shape 3** | (1)<br>0<br>(0.64) | (0.13)<br>2<br>(0.98) | (1)<br>0<br>(0.96) | (0.92)<br>3<br>(0.31) | 5 | (1)<br>0<br>(0.54) | (0.01)<br>4<br>(1) | (1)<br>0<br>(0.94) | (0.99)<br>3<br>(0.05) | 7 |
| **Shape 4** | (1)<br>0<br>(0.84) | (0.02)<br>2<br>(1) | (1)<br>0<br>(0.98) | (1)<br>0<br>(0.05) | 2 | (1)<br>0<br>(0.92) | (0.13)<br>1<br>(1) | (1)<br>0<br>(0.99) | (1)<br>0<br>(0.22) | 1 |
| **Shape 5** | (1)<br>0<br>(1) | (1)<br>0<br>(1) | (1)<br>0<br>(1) | (1)<br>0<br>(1) | 0 | (1)<br>0<br>(1) | (1)<br>0<br>(1) | (1)<br>0<br>(1) | (1)<br>0<br>(1) | 0 |
| **Shape 6** | (1)<br>0<br>(1) | (1)<br>0<br>(1) | (1)<br>0<br>(1) | (1)<br>0<br>(1) | 0 | (1)<br>0<br>(1) | (1)<br>0<br>(1) | (1)<br>0<br>(1) | (1)<br>0<br>(1) | 0 |
| **Shape 7** | (1)<br>196<br>(1.70E-09) | (3.96E-35)<br>646<br>(1) | (3.58E-09)<br>57<br>(1) | (1)<br>2,373<br>(2.75E-15) | 3,272 | (1)<br>167<br>(2.62E-14) | (3.76E-33)<br>622<br>(1) | (7.22E-06)<br>48<br>(1) | (1)<br>2,323<br>(2.57E-10) | 3,160 |
| **Shape 8** | (1.17E-40)<br>819<br>(1) | (1)<br>198<br>(5.71E-265) | (1)<br>3<br>(6.54E-27) | (7.56E-81)<br>5,791<br>(1) | 6,811 | (7.00E-36)<br>775<br>(1) | (1)<br>197<br>(7.66E-240) | (1)<br>2<br>(1.47E-26) | (2.93E-74)<br>5,513<br>(1) | 6,487 |
| **Total** | 1,443 | 2,196 | 140 | 13,199 | 16,978 | 1,443 | 2,196 | 140 | 13,199 | 16,978 |

**S4 Table.** Number of genes assigned to each distribution shape in time series datasets. Numbers in parentheses represent the P-values for enrichment (above) and depletion (below). The categories include differentially expressed (DE) and regulator (REG) genes – refer to methods for specific categories in Drosophila dataset.

|  | Drosophila Embryogenesis Dataset |  |  |  |  |  |  | Duck Preadipocyte Dataset |  |  |  |  |
| --- | --- | --- | --- | --- | --- | --- | --- | --- | --- | --- | --- | --- |
|  | Down/down | Down/up | Up/down | Up/up | REG | Others | Total | REG | DE | DE-REG | Others | Total |
| **Shape 1** | (4.29E-16)<br>292<br>(1) | (0.94)<br>305<br>(0.07) | (0.01)<br>76<br>(0.99) | (8.74E-04)<br>122<br>(1) | (1)<br>106<br>(4.17E-05) | (1)<br>638<br>(8.45E-13) | 1,433 | (0.97)<br>63<br>(0.05) | (1.17E-10)<br>584<br>(1) | (0.15)<br>29<br>(0.90) | (1)<br>1,307<br>(1.47E-08) | 1,983 |
| **Shape 2** | (0.16)<br>17<br>(0.90) | (0.96)<br>16<br>(0.08) | (0.02)<br>9<br>(0.99) | (0.63)<br>6<br>(0.53) | (0.98)<br>5<br>(0.06) | (0.69)<br>50<br>(0.39) | 98 | (0.43)<br>11<br>(0.69) | (1)<br>21<br>(1.71E-11) | (0.83)<br>2<br>(0.39) | (6.66E-10)<br>228<br>(1) | 262 |
| **Shape 3** | (0.14)<br>3<br>(0.96) | (0.41)<br>3<br>(0.83) | (1)<br>0<br>(0.65) | (1)<br>0<br>(0.51) | (1)<br>0<br>(0.34) | (0.87)<br>4<br>(0.31) | 10 | 1<br>0<br>(0.85) | 1<br>0<br>(0.34) | 1<br>0<br>(0.95) | (0.26)<br>4<br>1 | 4 |
| **Shape 4** | (1)<br>0<br>(1) | (1)<br>0<br>(1) | (1)<br>0<br>(1) | (1)<br>0<br>(1) | (1)<br>0<br>(1) | (1)<br>0<br>(1) | 0 | (1)<br>0<br>(1) | (1)<br>0<br>(1) | (1)<br>0<br>(1) | (1)<br>0<br>(1) | 0 |
| **Shape 5** | (1)<br>0<br>(1) | (1)<br>0<br>(1) | (1)<br>0<br>(1) | (1)<br>0<br>(1) | (1)<br>0<br>(1) | (1)<br>0<br>(1) | 0 | (1)<br>0<br>(1) | (1)<br>0<br>(1) | (1)<br>0<br>(1) | (1)<br>0<br>(1) | 0 |
| **Shape 6** | (1)<br>468<br>(8.76E-17) | (1.38E-18)<br>1,159<br>(1) | (0.02)<br>202<br>(0.98) | (1)<br>210<br>(3.46E-13) | (0.07)<br>467<br>(0.94) | (0.12)<br>2,351<br>(0.89) | 4,390 | (0.58)<br>54<br>(0.47) | (0.83)<br>324<br>(0.18) | (0.06)<br>24<br>(0.97) | (0.32)<br>1,023<br>(0.71) | 1,425 |
| **Shape 7** | (1)<br>1<br>(3.27E-51) | (1)<br>16<br>(1.6E-69) | (0.20)<br>38<br>(0.84) | (4.75E-51)<br>171<br>(1) | (2.62E-05)<br>115<br>(1) | (6.07E-29)<br>564<br>(1) | 790 | (0.90)<br>117<br>(0.12) | (5.42E-11)<br>930<br>(1) | (9.71E-04)<br>58<br>(1) | (1)<br>2,227<br>(1.06E-10) | 3,332 |
| **Shape 8** | (1.43E-29)<br>267<br>(1) | (0.01)<br>271<br>(0.99) | (1)<br>1<br>(5.10E-20) | (1)<br>2<br>(1.48E-30) | (0.85)<br>98<br>(0.18) | (1)<br>509<br>(9.72E-04) | 1,050 | (0.01)<br>270<br>(0.99) | (1)<br>1,302<br>(5.3E-16) | (1)<br>47<br>(2.2E-06) | (1.05E-14)<br>4,697<br>(1) | 6,316 |
| **Total** | 1,048 | 1,770 | 326 | 511 | 791 | 4,116 | 7,771 | 515 | 3,161 | 160 | 9,486 | 13,322 |

**S5 Table.** Number of genes assigned to each distribution shape in human dataset. Numbers in parentheses represent the P-values for enrichment (above) and depletion (below). The categories include differentially expressed (DE), regulator (REG) and tissue enriched (TE) genes.

|  | Human Dataset |  |  |  |  | Total |
| --- | --- | --- | --- | --- | --- | --- |
|  | REG | DE | DE-REG | TE | Others |  |
| **Shape 1** | (0.55)<br>215<br>(0.48) | (2.48E-04)<br>182<br>(1) | (0.02)<br>24<br>(0.99) | (0.09)<br>55<br>(0.93) | (1)<br>3,411<br>(1.53E-03) | 3,881 |
| **Shape 2** | (9.31E-06)<br>344<br>(1) | (3.28E-10)<br>262<br>(1) | (2.92E-04)<br>35<br>(1) | (0.99)<br>46<br>(0.01) | (1)<br>4,382<br>(6.34E-13) | 5,065 |
| **Shape 3** | (0.92)<br>1<br>(0.30) | (4.82E-03)<br>6<br>(1) | (0.16)<br>1<br>(0.99) | (1)<br>0<br>(0.60) | (0.96)<br>35<br>(0.09) | 43 |
| **Shape 4** | (0.37)<br>1<br>(0.93) | (1)<br>0<br>(0.74) | (1)<br>0<br>(0.97) | (1)<br>0<br>(0.91) | (0.79)<br>7<br>(0.60) | 8 |
| **Shape 5** | (1)<br>0<br>(1) | (1)<br>0<br>(1) | (1)<br>0<br>(1) | (1)<br>0<br>(1) | (1)<br>0<br>(1) | 0 |
| **Shape 6** | (1)<br>0<br>(1) | (1)<br>0<br>(1) | (1)<br>0<br>(1) | (1)<br>0<br>(1) | (1)<br>0<br>(1) | 0 |
| **Shape 7** | (1)<br>79<br>(4.00E-03) | (1)<br>48<br>(2.94E-03) | (0.88)<br>5<br>(0.22) | (7.21E-34)<br>91<br>(1) | (9.40E-01)<br>1,640<br>(0.07) | 1,860 |
| **Shape 8** | (0.99)<br>435<br>(0.01) | (1)<br>217<br>(5.79E-14) | (1)<br>13<br>(3.59E-07) | (1)<br>39<br>(2.65E-18) | (3.58E-24)<br>7,733<br>(1) | 8,427 |
| **Total** | 1,075 | 715 | 78 | 231 | 17,208 | 19,284 |

**S6 Table.** Number of genes assigned to each distribution shape in all datasets based on genes being on the top or bottom 5% when ranked by degree (number of significant correlations to other genes). Numbers in parentheses represent the P-values for enrichment (above) and depletion (below).

|  | **Feed Efficiency** |  | **Puberty** |  | **Drosophila** |  | **Duck** |  | **Human** |  |
| --- | --- | --- | --- | --- | --- | --- | --- | --- | --- | --- |
|  | **Top** | **Bottom** | **Top** | **Bottom** | **Top** | **Bottom** | **Top** | **Bottom** | **Top** | **Bottom** |
| **Shape 1** | (1)<br>0<br>(8.44E-50) | (4.72E-24)<br>198<br>(1) | (1)<br>0<br>(3.27E-194) | (1.56E-37)<br>518<br>(1) | (1)<br>0<br>(3.74E-36) | (7.1E-126)<br>280<br>(1) | (1)<br>0<br>(1.18E-48) | (9.3E-110)<br>335<br>(1) | (1)<br>0<br>(9.50E-98) | (1)<br>61<br>(3.78E-35) |
| **Shape 2** | (1)<br>0<br>(8.41E-20) | (4.27E-252)<br>335<br>(1) | (1)<br>0<br>(1.93E-13) | (8.64E-275)<br>314<br>(1) | (1)<br>0<br>(6.31E-03) | (8.84E-44)<br>53<br>(1) | (1)<br>0<br>(1.27E-06) | (2.3E-280)<br>231<br>(1) | (1)<br>0<br>(1.90E-132) | (<1E-999)<br>894<br>(1) |
| **Shape 3** | (1)<br>5<br>(1.54E-12) | (1)<br>18<br>(6.76E-05) | (1)<br>0<br>(0.90) | (2.50E-03)<br>2<br>(1) | (1)<br>0<br>(5.98E-01) | (2.68E-06)<br>6<br>(1) | (1)<br>0<br>(0.81) | (6.19E-06)<br>4<br>(1) | (1)<br>0<br>(0.11) | (4.02E-05)<br>10<br>(1) |
| **Shape 4** | (1.38E-94)<br>578<br>(1) | (1)<br>31<br>(6.97E-233) | (1)<br>0<br>(0.90) | (2.50E-03)<br>2<br>(1) | (1)<br>0<br>(1) | (1)<br>0<br>(1) | (1)<br>0<br>(1) | (1)<br>0<br>(1) | (1)<br>0<br>(0.66) | (0.34)<br>1<br>(0.94) |
| **Shape 5** | (1)<br>0<br>(1) | (1)<br>0<br>(1) | (1)<br>0<br>(1) | (1)<br>0<br>(1) | (1)<br>0<br>(1) | (1)<br>0<br>(1) | (1)<br>0<br>(1) | (1)<br>0<br>(1) | (1)<br>0<br>(1) | (1)<br>0<br>(1) |
| **Shape 6** | (1)<br>0<br>(1) | (1)<br>0<br>(1) | (1)<br>0<br>(1) | (1)<br>0<br>(1) | (1.31E-100)<br>389<br>(1) | (1)<br>27<br>(3.3E-101) | (1)<br>0<br>(2.41E-34) | (1)<br>1<br>(2.06E-32) | (1)<br>0<br>(1) | (1)<br>0<br>(1) |
| **Shape 7** | (1)<br>0<br>(8.32E-03) | (0.99)<br>1<br>(0.05) | (1.54E-31)<br>299<br>(1) | (1)<br>5<br>(5.83E-70) | (1)<br>0<br>(2.48E-19) | (1)<br>13<br>(1.99E-07) | (0.99)<br>142<br>(0.01) | (1)<br>45<br>(2.35E-36) | (1)<br>1<br>(2.13E-42) | (1)<br>0<br>(1.94E-44) |
| **Shape 8** | (1)<br>0<br>(9.45E-04) | (1)<br>0<br>(9.45E-04) | (1.77E-58)<br>550<br>(1) | (1)<br>8<br>(3.03E-166) | (1)<br>0<br>(6.20E-26) | (1)<br>10<br>(2.14E-14) | (1.31E-64)<br>524<br>(1) | (1)<br>50<br>(8.6E-118) | (<1E-999)<br>965<br>(1) | (1)<br>0<br>(2.81E-250) |
| **Total** | 583 | 583 | 849 | 849 | 389 | 389 | 666 | 666 | 966 | 966 |

## References

1. Swami M. Networking complex traits. Nat Rev Genet. 2009;10: 2566. doi:10.1038/ng.332

2. Hudson NJ, Dalrymple BP, Reverter A. Beyond differential expression: the quest for causal mutations and effector molecules. BMC Genomics. 2012. doi:10.1186/1471-2164-13-356

3. Mar JC, Matigian NA, Mackay-Sim A, Mellick GD, Sue CM, Silburn PA, et al. Variance of gene expression identifies altered network constraints in neurological disease. Gibson G, editor. PLoS Genet. 2011;7: e1002207. doi:10.1371/journal.pgen.1002207

4. Barabási A-L, Oltvai ZN. Network biology: understanding the cell’s functional organization. Nat Rev Genet. 2004;5: 101–13. doi:10.1038/nrg1272

5. Hudson NJ, Reverter A, Wang YH, Greenwood PL, Dalrymple BP. Inferring the transcriptional landscape of bovine skeletal muscle by integrating co-expression networks. PLoS One. 2009;4. doi:10.1371/journal.pone.0007249

6. Remondini D, O’Connell B, Intrator N, Sedivy JM, Neretti N, Castellani GC, et al. Targeting c-Myc-activated genes with a correlation method: Detection of global changes in large gene expression network dynamics. Proc Natl Acad Sci. 2005;102: 6902–6906. doi:10.1073/pnas.0502081102

7. R Core Team. R: A language and environment for statistical computing. R Foundation for Statistical Computing. Vienna; 2019. Available: https://www.r-project.org/

8. Reverter A, Chan EKF. Combining partial correlation and an information theory approach to the reversed engineering of gene co-expression networks. Bioinformatics. 2008;24: 2491–2497. doi:10.1093/bioinformatics/btn482

9. Wang Z, Yin Z-T, Zhang F, Li X-Q, Chen S-R, Yang N, et al. Dynamics of transcriptome changes during subcutaneous preadipocyte differentiation in ducks. BMC Genomics. 2019. doi:10.1186/s12864-019-6055-9

10. Lonsdale J, Thomas J, Salvatore M, Phillips R, Lo E, Shad S, et al. The Genotype-Tissue Expression (GTEx) project. Nat Genet. 2013;45: 580–585. doi:10.1038/ng.2653

11. Alexandre PA, Kogelman LJA, Santana MHA, Passarelli D, Pulz LH, Fantinato-Neto P, et al. Liver transcriptomic networks reveal main biological processes associated with feed efficiency in beef cattle. BMC Genomics. 2015;16. doi:10.1186/s12864-015-2292-8

12. Alexandre PA, Naval-Sanchez M, Porto-Neto LR, Ferraz JBS, Reverter A, Fukumasu H. Systems biology reveals NR2F6 and TGFB1 as key regulators of feed efficiency in beef cattle. Front Genet. 2019;10: 1–16. doi:10.3389/fgene.2019.00230

13. Koch RM, Swiger LA, Chambers D, Gregory KE. Efficiency of Feed Use in Beef Cattle. J Anim Sci. 1963;22: 486–494.

14. Zhang HM, Liu T, Liu CJ, Song S, Zhang X, Liu W, et al. AnimalTFDB 2.0: A resource for expression, prediction and functional study of animal transcription factors. Nucleic Acids Res. 2015;43: D76–D81. doi:10.1093/nar/gku887

15. Nguyen LT, Reverter A, Cánovas A, Venus B, Anderson ST, Islas-Trejo A, et al. STAT6, PBX2, and PBRM1 Emerge as Predicted Regulators of 452 Differentially Expressed Genes Associated With Puberty in Brahman Heifers. Front Genet. 2018;9: 87. doi:10.3389/fgene.2018.00087

16. Fortes MRS, Zacchi LF, Nguyen LT, Raidan F, Weller MMDCA, Choo JJY, et al. Pre- and post-puberty expression of genes and proteins in the uterus of *Bos indicus* heifers: the luteal phase effect post-puberty. Anim Genet. 2018;49: 539–549. doi:10.1111/age.12721

17. Nguyen LT, Reverter A, Cánovas A, Venus B, Islas-Trejo A, Porto-Neto LR, et al. Global differential gene expression in the pituitary gland and the ovaries of pre- and postpubertal Brahman heifers1. J Anim Sci. 2017;95: 599–615. doi:10.2527/jas.2016.0921

18. Fortes MRS, Nguyen LT, Weller MMDCA, Cánovas A, Islas-Trejo A, Porto-Neto LR, et al. Transcriptome analyses identify five transcription factors differentially expressed in the hypothalamus of post-versus prepubertal Brahman heifers. J Anim Sci. 2016;94: 3693–3702. doi:10.2527/jas.2016-0471

19. Hu H, Miao YR, Jia LH, Yu QY, Zhang Q, Guo AY. AnimalTFDB 3.0: A comprehensive resource for annotation and prediction of animal transcription factors. Nucleic Acids Res. 2019. doi:10.1093/nar/gky822

20. Becker K, Bluhm A, Casas-Vila N, Dinges N, Dejung M, Sayols S, et al. Quantifying post-transcriptional regulation in the development of Drosophila melanogaster. Nat Commun. 2018;9: 4970. doi:10.1038/s41467-018-07455-9

21. Rhee DY, Cho DY, Zhai B, Slattery M, Ma L, Mintseris J, et al. Transcription factor networks in Drosophila melanogaster. Cell Rep. 2014;8: 2031–2043. doi:10.1016/j.celrep.2014.08.038

22. Uhlen M, Fagerberg L, Hallstrom BM, Lindskog C, Oksvold P, Mardinoglu A, et al. Tissue-based map of the human proteome. Science (80-). 2015;347: 1260419–1260419. doi:10.1126/science.1260419

23. Crow M, Lim N, Ballouz S, Pavlidis P, Gillis J. Predictability of human differential gene expression. Proc Natl Acad Sci. 2019;116: 6491–6500. doi:10.1073/pnas.1802973116

24. Lee HK, Hsu AK, Sajdak J, Qin J, Pavlidis P. Coexpression Analysis of Human Genes Across Many Microarray Data Sets. Genome Res. 2004;14: 1085–1094. doi:10.1101/gr.1910904

25. Reverter A, Hudson NJ, Nagaraj SH, Pérez-Enciso M, Dalrymple BP. Regulatory impact factors: Unraveling the transcriptional regulation of complex traits from expression data. Bioinformatics. 2010;26: 896–904. doi:10.1093/bioinformatics/btq051

26. Cánovas A, Reverter A, DeAtley KL, Ashley RL, Colgrave ML, Fortes MRS, et al. Multi-Tissue Omics Analyses Reveal Molecular Regulatory Networks for Puberty in Composite Beef Cattle. PLoS One. 2014;9: e102551. doi:10.1371/journal.pone.0102551

